# Strategic Segmentation: A Nash Equilibrium based Approach for Weed Segmentation in Agricultural Fields

**DOI:** 10.64898/2026.01.27.702114

**Authors:** Shreyan Kundu, Souradeep Mukhopadhyay, Trishit Mukherjee, Shubhankar Mondal, Biswadip Basu Mallik

## Abstract

Weeds present a major challenge to agricultural productivity, competing with crops for critical resources like water, nutrients, and sunlight, resulting in significant yield reductions. Prompt weed identification is essential for enabling effective control strategies, such as the application of herbicides or mechanical removal, to minimize their impact on crop growth. This research focuses on developing a deep learning approach based on game theory for detecting weeds. Using CWFID dataset captured at various times and days, along with multispectral data in the visible and near-infrared spectrum, the study aims to improve early detection methods for more efficient weed management in agricultural settings. A novel segmentation technique for weed regions is introduced, employing a zero-sum game theory model to reconcile conflicting classifications from different weed detectors. These regions are treated as zones of conflict between weeds and crops, with each detector representing a different strategy. By defining an appropriate utility function, the method identifies the Nash equilibrium, effectively minimizing false positive detections of weeds.

## 1 Introduction

Weed detection is crucial for improving agricultural productivity and sustainability, as weeds compete with crops for water and nutrients. Manual weeding is impractical for large-scale farming due to labor costs, and robotic weeding has drawbacks such as energy consumption and crop damage. Herbicide use, common since the 1940s, faces challenges from weed mutations and legal restrictions, making precise weed identification essential for effective crop protection. This study introduces a novel weed segmentation approach by leveraging differences between multiple weed detectors rather than merging their outputs. The method formulates detector disagreements as a strictly competitive game, where weed and non-weed regions act as opposing players. Each classifier recognizing a region as weed defines a weed strategy, while other classifiers form non-weed strategies. A correlation distance function ensures homogeneity and adapts dynamically using weed samples from the image. The proposed method significantly reduces false positives by treating the non-weed player as a decision agent.

(1) A novel fusion method enhancing weed segmentation by analyzing classifier disagreements.
(2) A zero-sum game model for effective weed and non-weed interaction modeling.
(3) A utility function reducing false positives by incorporating region distances and correlation with weed samples, improving segmentation accuracy.

Experimental results validate the effectiveness and adaptability of this approach in binary classification tasks requiring multiple classifiers.

## 2 Related Work

Distinguishing crops from weeds in agricultural fields presents a significant challenge due to their similar visual features. Traditional segmentation methods typically involve pre-processing followed by pixel-based classification[1, 2]. Pre-processing techniques aim to reduce noise and mitigate illumination effects through image enhancement, while pixel-based segmentation relies on methods such as color index-based, threshold-based, and machine learning-based approaches. In addition, multi-modal approaches that combine RGB cameras with hyperspectral imaging have proven effective in addressing the variations in crop and weed appearances, thus enhancing segmentation accuracy. By capturing a broader range of spectral information, these methods can identify subtle differences between weeds and crops that are not easily detectable with conventional RGB imaging. Hamuda provides a detailed review of image processing techniques for plant extraction and segmentation in agricultural fields. Their review covers a wide range of methods, providing insights into the effectiveness of different techniques. However, it lacks a detailed performance analysis under varying environmental conditions, which is crucial for real-world applications. Lottes et al. [3] presents a classification system for distinguishing sugar beets from weeds, demonstrating its potential for precision farming. While effective, the method faces challenges in scalability and adaptability across diverse field conditions. Further validation is needed to ensure robustness in varying agricultural settings.Confidence-aware ensemble fusion has also been explored in medical image analysis, where Kundu et al. [26] introduced weighted confidence-based ensembling with attention to improve robust decision-making, aligning with our conflict-aware fusion strategy. Edge-aware and multi-scale feature integration has been shown to improve segmentation robustness, as demonstrated by Kar et al. [16], motivating structured fusion and boundary-sensitive decision modeling in our framework. Attention-guided feature fusion has also been applied to plant disease analysis; Kundu et al. [19] employed ensemble CNNs with attention for potato leaf disease classification, reinforcing the effectiveness of complementary model fusion in agricultural image understanding, this aided us for the plant health understanding.

Guerrero et al. [4] apply SVMs to crop and weed classification in maize fields, highlighting the potential of machine learning in agriculture. Despite its high accuracy, feature selection and model generalization remain challenges. Kundu et al. [24] employed majority-voting ensembling with contrastive learning for malaria cell detection, motivating ensemble-driven fusion strategies for robust visual decision-making. Milioto et al. [5] propose a real-time semantic segmentation method using CNNs for crop and weed detection. Although promising for autonomous agricultural robots, computational complexity and model optimization are potential barriers to real-world deployment. Economy-inspired feature selection and fusion strategies have been shown to enhance discriminative learning in medical imaging; Kundu et al. [11] employed Gini-based feature selection with linear production–inspired fusion, motivating utility-driven and structured fusion mechanisms in our framework. You et al. [6] propose a DNN-based semantic segmentation method for weed and crop detection, achieving high detection accuracy. However, challenges such as dataset diversity and the computational resources required for model training must be addressed for practical implementation. Kundu et al. [25] demonstrated multi-CNN feature fusion for rice crop disorder detection, motivating complementary model fusion for robust agricultural image segmentation. Fuzzy and rank-based ensemble decision strategies have also been explored in biomedical imaging; Kundu et al. [27] combined fuzzy ranking with contrastive learning for malaria detection, motivating confidence-aware and structured fusion mechanisms in our framework.

Intawong et al. [7] introduced a pixel-based quality measure for segmentation algorithms, integrating precision, recall, and specificity to enhance performance evaluation. Kouzana, Nakib, and Dogaz [9] applied Branch and Bound algorithms with Game Theory for microscopic image segmentation, improving accuracy and efficiency, with potential applications in medical and biological image analysis. Hybrid supervised–contrastive learning has shown promise in agricultural vision. Kundu et al. [22] demonstrated improved feature discrimination for jute pest classification, motivating disagreement-aware decision modeling for fine-grained weed segmentation. Attention-enhanced feature fusion has been explored for histopathology-based cancer grading. Kundu et al. [23] proposed RAF2Net, which fuses multiple lightweight CNN backbones with attention, motivating structured fusion strategies that exploit complementary model responses, as adopted in our segmentation framework.

## 3 Our Proposed Algorithm

Identifying weeds is a binary classification task, distinguishing between weed and non-weed pixels. Conflicts, or ‘conflict areas’, arise when different weed detection methods, based on cues like color or texture, yield opposing results. This paper models these conflicts as an evolutionary game between two players: the weed player and the non-weed player, each striving to dominate regions aligned with their nature. Framed as a strictly competitive game (null sum game), first player’s gain is the other’s loss and vice versa. The game’s mixed saddle point serves as a compromise, reducing false positives by determining territory allocation based on the persuasiveness of detection strategies.

### 3.1 Designing the zero-sum game

Let 𝒲 be the set of weed classifiers. The zero-sum game 𝒢 = (𝒥, 𝒲_*t*_, 𝒰) is defined as follows where, 𝒥 = {*weedplayer, non* − *weedplayer*}.

Let 𝒲_*t*_ (for *t* = 1, 2) represent the sets of pure strategies for each player in a conflict patch 𝒫, where 𝒲_*t*_ are subsets of the overall strategy set 𝒲.

- *W*_1_ = {𝒲_*t*_ ∈ 𝒲 : 𝒲_*t*_ classified 𝒫 as weed}.
- *W*_2_ = 𝒲\*W*_1_.

The weed player’s strategies include classifiers that label 𝒫 as weed, and the non-weed player’s strategies consist of the remaining classifiers.

### 3.2 Description of Utility Function 𝒰

The key focus of this research is defining the payoff function. Based on human visual perception, homogeneity and similarity are used to establish the function. To determine if the conflict region resembles weed, we define a weed-correlation distance 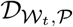, reflecting visual similarity to weeds. The distance is calculated for a conflict patch 𝒫 by comparing it with regions classified as weed by all classifiers in 𝒲, ensuring uniformity and resemblance. For each strategy 𝒲_*t*_ ∈ 𝒲, we define a weed data matrix 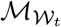, where each row represents the weed feature vector of 𝒲_*t*_ from a patch classified as weed by all strategies 𝒲_*t*_ ∈ 𝒲. For instance, the feature vector corresponding to the RGB strategy consists of the *R, G*, and *B* channel components of the weed.

Let take Δ_*j*_ and 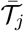 as the standard deviation and average of the *j*-th variable in 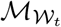, given by:

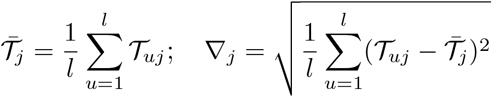

where *l* is the number of weed patches and *n* is the dimension of the feature vector. The center of gravity of 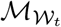 is:

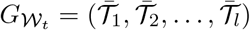

The S.D of 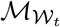 is:

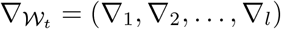

The centered and standardized matrix 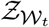 is derived as:

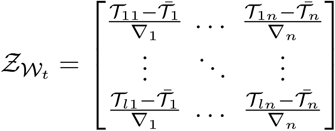

The correlation matrix 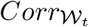 is then computed as:

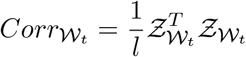

Let 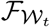 be the feature vector of 𝒫 for strategy 𝒲_*t*_:

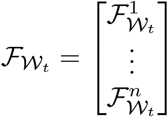

The centered and standardized vector 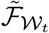 is:

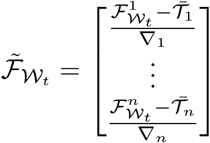

Let 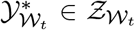 minimize the Euclidean distance 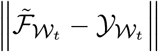. The vector 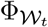 is defined as:

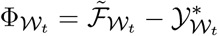

The correlation distance 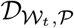, quantifying the similarity between patch 𝒫 and the weed matrix 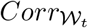, is:

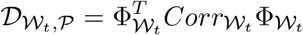

This distance approaches zero when 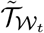 resembles weed features, indicating similarity to weed regions.

To compare classifiers *W*_1_ (weed) and *W*_2_ (non-weed), the utility function 𝒰 (*W*_1_, *W*_2_) quantifies the similitude of patch 𝒫 to weed pixels classified by 𝒲_1_ and its dissimilarity according to 𝒲_2_. This is done through homogeneity norms:

Similarity *λ* to weed pixels is:

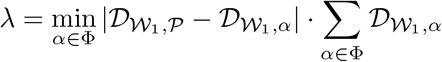

Dissimilarity *β* to non-weed pixels is:

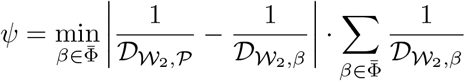

Where 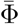 represents the set of non-weed neighbors of 𝒫 according to the classification of strategy 𝒲_2_.

The payoff function is:

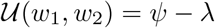

This reflects the interplay between the weed and non-weed strategies. 𝒰 (*w*_1_, *w*_2_) > 0 implies 𝒫 is more similar to weed, while 𝒰 (*w*_1_, *w*_2_) < 0 implies non-weed similarity.

### 3.3 Computation of Nash Equilibrium

In the framework of game theory, the categorization between weed and non-weed classes can be resolved by computing the saddle point or Nash Equilibrium for a contending area. The class assignment for a zero-sum game is determined by the game’s value, which is equal to the payoff value obtained from each player’s optimum tactics. Fairness is indicated by a value of zero, whereas the weed class is favored by a positive value and the non-weed class is favored by a negative value.

When the pure game lacks a definite value, we extend it to a mixed strategy game, denoted by 𝒢^*^, characterized as 𝒢^*^ = (𝒥, Δ(𝒲_*t*_), 𝒰), where Δ(𝒲_*t*_) represents the probability distributions over 𝒲_*t*_:

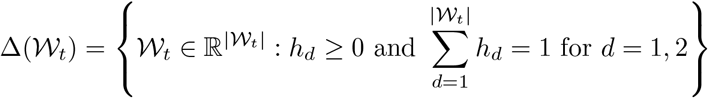

Here, *υ* = *E*(𝒰 (*i, j*)) is the expected payoff, calculated from the payoff function 𝒰 (*i, j*) for strategies (*i, j*) ∈ Δ(𝒲_1_) × Δ(𝒲_2_), represented as:

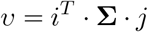

where Σ is the game’s payoff matrix.

Von Neumann’s Minimax Theorem ensures the existence of a unique value *υ*^*^ in mixed finite games, defined as:

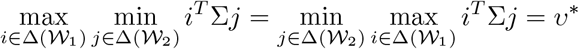

with optimal mixed strategies (*i*^*^, *j*^*^), and the expected payoff calculated as:

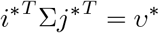

Thus, *υ*^*^ represents Player 1’s maximum floor and Player 2’s minimum ceiling. To solve for (*i*^*^, *j*^*^, *υ*^*^), a LPP is formulated. For Weed-player, the problem is:

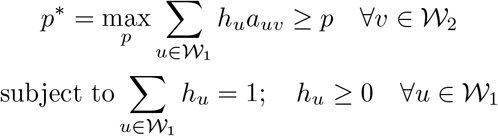

Similarly, for non-weed-player, the dual LP is:

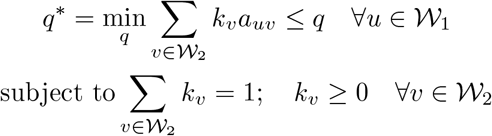

This study proposes a weed detection system integrating both rule-based and machine learning methods. Rule-based techniques isolate weed-colored pixels by thresholding RGB, HSV, and YCbCr color spaces, employing the Peer model in RGB and chrominance channels (Hue, Saturation, Cb, Cr) in HSV and YCbCr spaces. Machine learning approaches utilize Artificial Neural Networks (ANN) trained on texture and color features. Weed textures are analyzed via Gabor filters, generating 13 Haralick indices. ANN classifiers trained either solely on texture (Texture-ANN) or combined texture-color features (Color-Texture-ANN) enhance accuracy. A zero-sum game model resolves classifier conflicts, and parallel architecture ensures efficient processing of large images.

## 4 Results

### 4.1 Dataset Description

CWFID dataset includes 2000 grass samples for model training, with 1200 and 800 images from the grass weed species ‘Setaria verticillata’ and ‘Digitaria sanguinalis’ available online (http://github.com/cwfid). Each image has a Ground Truth vegetation segmentation mask and manual crop vs. weed plant category annotation.

### 4.2 Setup for Experiment

The 24-GB NVIDIA A5000 Tensor Core GPU and Python 3.9 programming language were used in every trial PC. Using the PyTorch package, the deep learning model is developed in a CUDA-enabled environment. The Apex package from NVIDIA is integrated for mixed precision training, enabling efficient use of memory and maximizing performance on the A5000 GPU. To further optimize model training and manage distributed computing, PyTorch Lightning is employed, ensuring streamlined handling of training loops and boosting overall model performance.

### 4.3 Outcomes and Visual Illustration

In this section, we summarize the performance metrics of a model evaluated on the test set. The test loss is 0.426, indicating the model’s error, and the accuracy is 93.8%, showing how well the model correctly classifies the data. The Dice Score (0.783) and IoU Score (0.643) reflect the overlap between predicted and actual labels, commonly used in segmentation tasks. The precision score (0.932) shows the model’s accuracy in identifying positive instances, while the recall score (0.673) indicates how well the model captures all relevant positives. Overall, the model performs well but could improve in recall and IoU. The Model accuracy and loss curve is shown in Fig. 1 and 2 and comparisons in Fig. 5 and 6.

**Fig. 1:**
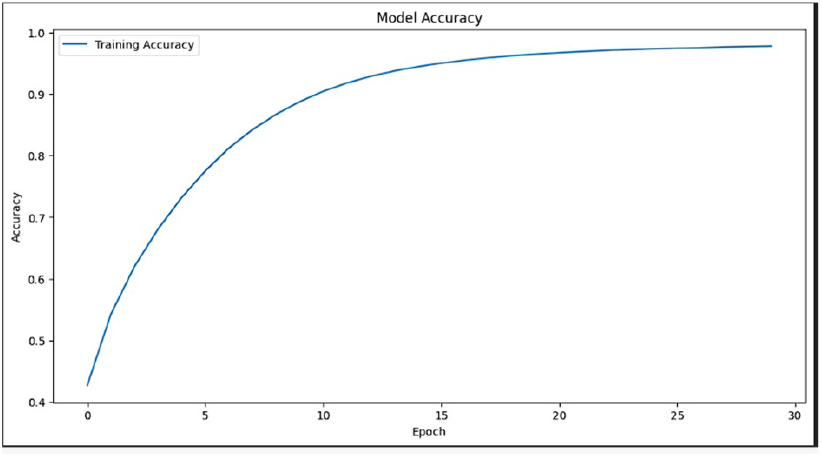
Model Accuracy Curve

**Fig. 2:**
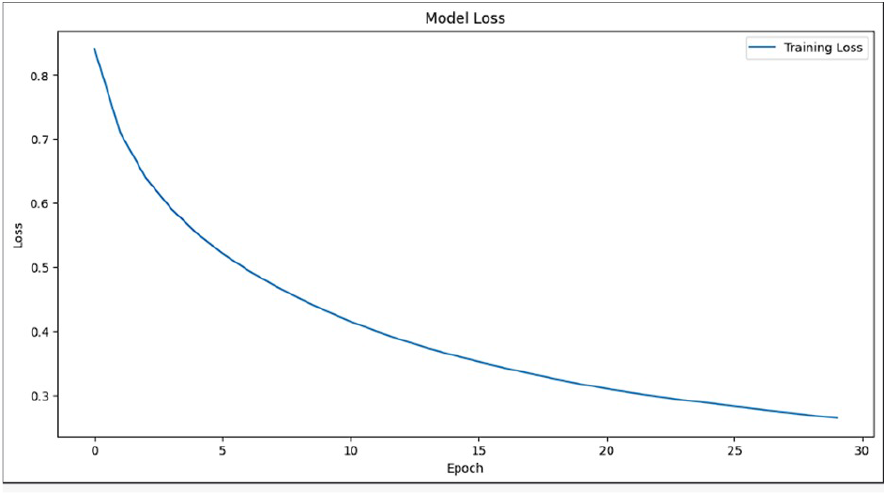
Model Loss Curve

The visual outputs are shown in Fig.3.

**Fig. 3:**
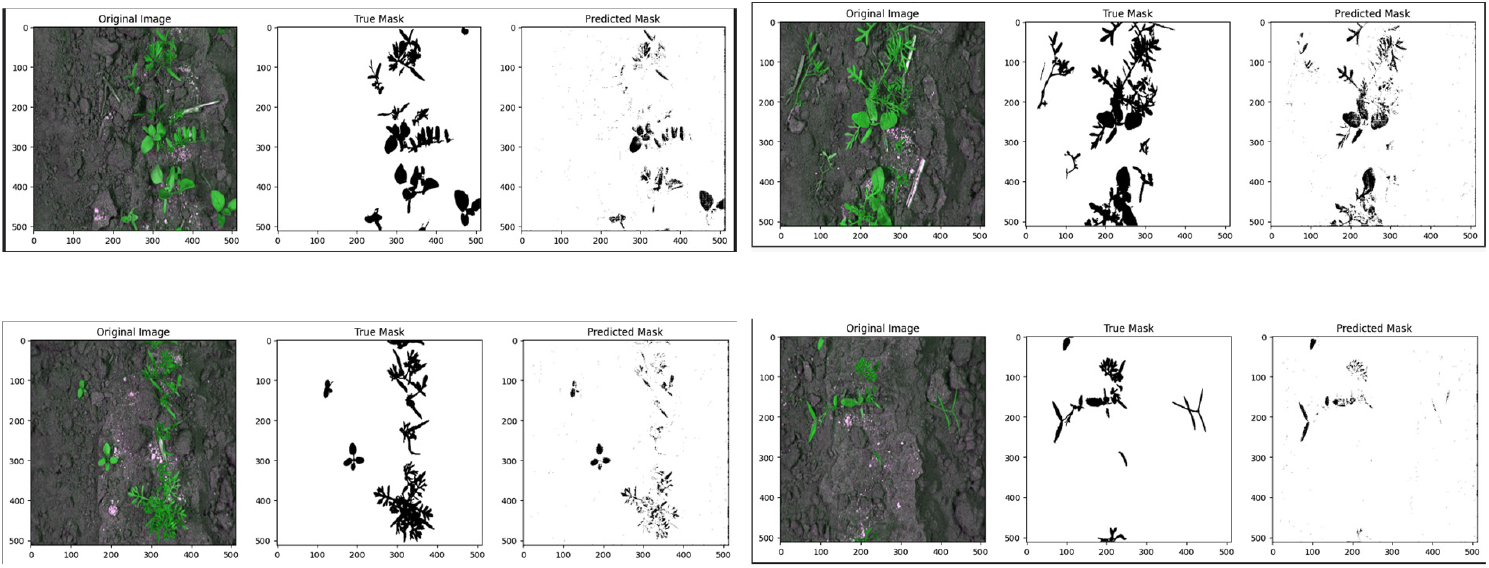
Visual outputs of Segmentation

Changes in Recall, Precision, Dice Score, and IOU score over Epochs are shown in Fig. 4.

**Fig. 4:**
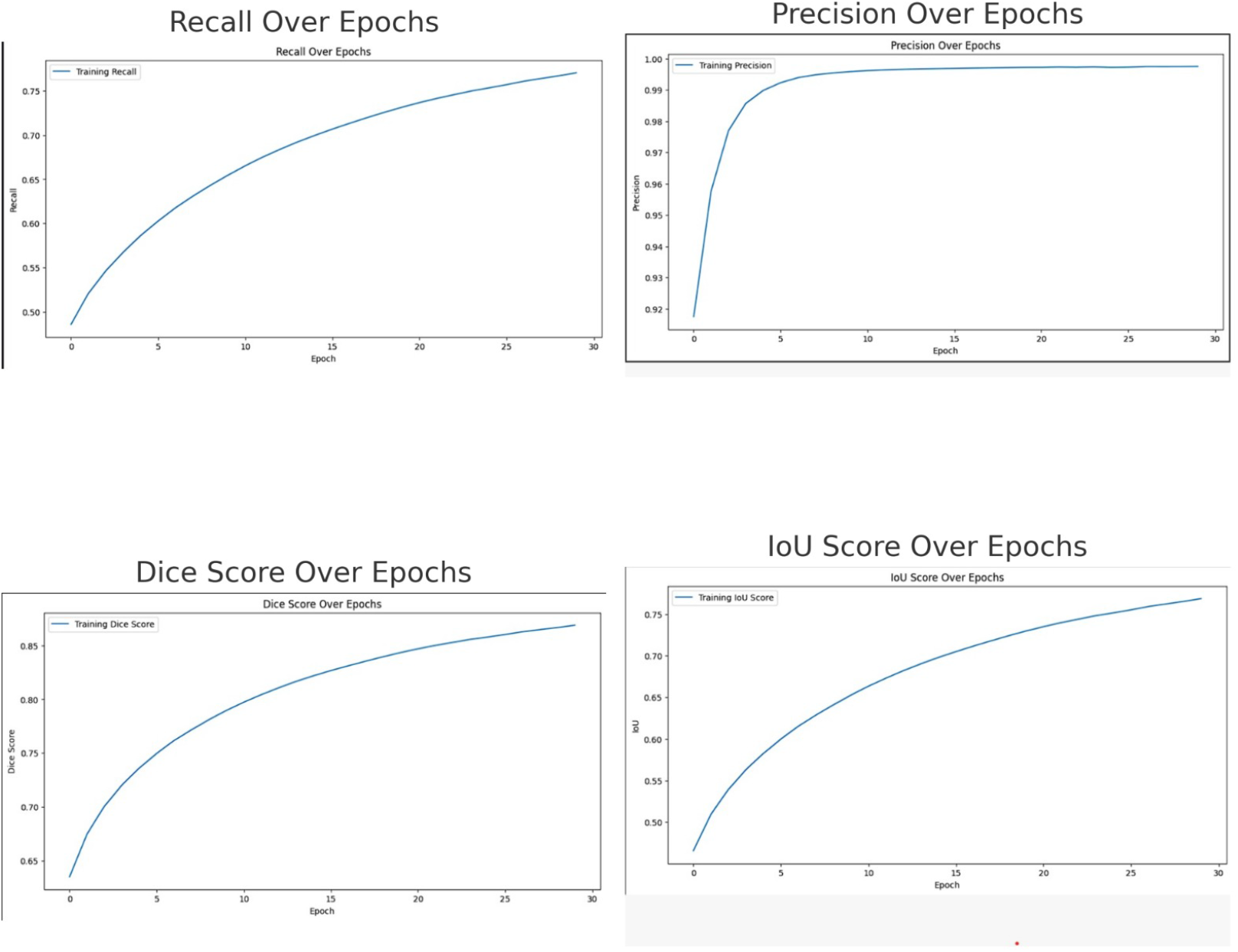
Changes in Recall, Precision, Dice Score, and IOU score over Epochs

**Fig. 5:**
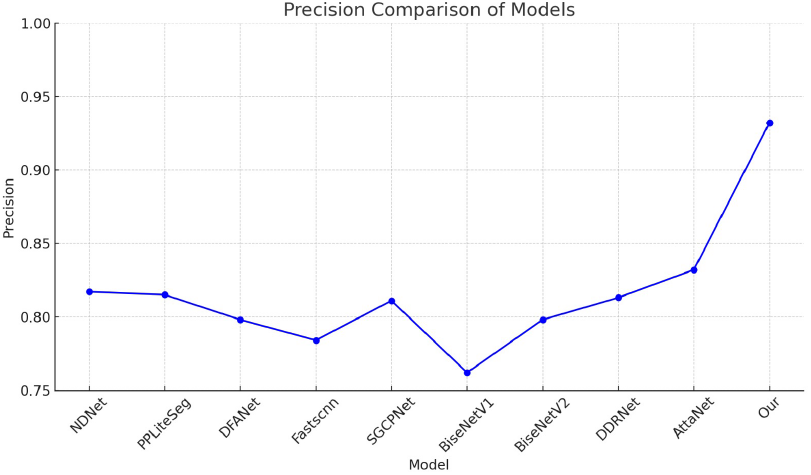
Precision Comparison

**Fig. 6:**
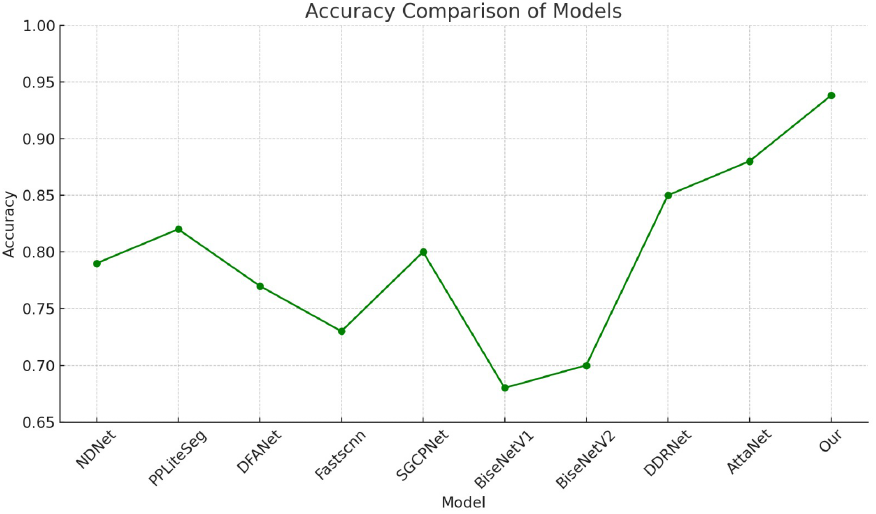
Accuracy Comparison

### 4.4 Comparison with other method

In the precision comparison, our method outperforms all other models with a precision of 0.932. AttaNet[10] follows at 0.832, while models like NDNet[12] (0.817), PPLiteSeg[13] (0.815), and DDRNet[14](0.813) also perform relatively well, albeit significantly lower than ours. Other models, such as DFANet[15] (0.798), BiseNetV2[18] (0.798), and SGCPNet[20] (0.811), are clustered around the 0.80 mark, with Fastscnn[21] (0.784) and BiseNetV1[17] (0.762) trailing behind. For accuracy, our model again leads with 0.938, ahead of AttaNet[10]’s 0.88 and DDRNet’s 0.85. Most other models, like SGCPNet[20] (0.80) and PPLiteSeg[13] (0.82), perform decently but remain below our model. DFANet[15] (0.77), BiseNetV2[18] (0.70), and BiseNetV1[17] (0.68) show noticeably lower accuracy. Fastscnn [21] lags with a 0.73 accuracy. Overall, our method demonstrates clear superiority across both precision and accuracy, particularly excelling over AttaNet and DDRNet, with a significant margin over other models.

## 5 Conclusion and future direction

The segmentation model performed well across multiple metrics, such as precision, recall, IoU, and Dice score. High precision (0.932) reflects its accuracy in identifying positive instances, while a recall of 0.673 suggests some difficulty in capturing all relevant positives. The IoU score (0.643) and Dice score (0.783) show an acceptable overlap between predicted and actual labels, which is crucial for effective segmentation. Despite these promising results, the recall and IoU scores highlight areas for further refinement, especially for tasks that demand comprehensive coverage of positive instances. Future work could focus on improving recall by experimenting with different loss functions, increasing the diversity of training datasets, and implementing advanced architectures, such as transformers or attention mechanisms. Additionally, incorporating domain adaptation techniques and unsupervised learning may enhance the model’s generalizability to new environments. Real-world applications could benefit from these improvements, particularly in complex tasks like weed detection or medical image segmentation. Future work will investigate integrating the proposed weed segmentation framework with IoT-based precision agriculture systems. Leveraging real-time soil and environmental sensing from autonomous rover–sensor networks, as demonstrated in IoT-enabled maize cultivation systems [8], can enable context-aware weed management and data-driven crop intervention strategies. Edge-enabled multi-agent perception frameworks have demonstrated the feasibility of low-latency, distributed visual decision-making; Mukherjee et al. [28] showed effective edge-assisted UAV–robot coordination, motivating future integration of our conflict-aware weed segmentation with decentralized, real-time agricultural robotic systems.

## References

[1] Pramanik, P., Mukhopadhyay, S., Mirjalili, S., & Sarkar, R. (2022). Deep feature selection using local search embedded social ski-driver optimization algorithm for breast cancer detection in mammograms. Neural Computing & Applications, 35, 5479–5499.

[2] Pramanik, P., Mukhopadhyay, S., Kaplun, D., Sarkar, R. (2022). A Deep Feature Selection Method for Tumor Classification in Breast Ultrasound Images. In: Tchernykh, A., Alikhanov, A., Babenko, M., Samoylenko, I. (eds) Mathematics and its Applications in New Computer Systems. MANCS 2021. Lecture Notes in Networks and Systems, vol 424. Springer, Cham.

[3] Lottes, P., Behley, J., Chebrolu, N., & Stachniss, C. (2016). A classification system for distinguishing sugar beets from weeds in precision farming. In Proceedings of ICRA 2016, Stockholm, Sweden.

[4] Guerrero, J. M., Pajares, G., Montalvo, M., Romeo, J., Guijarro, M., & Cruz, J. M. (2012). Support vector machines for crop and weed identification in maize fields. Expert Systems with Applications, 39(12), 11168–11175.

[5] Milioto, A., Lottes, P., & Stachniss, C. (2018). Real-time semantic segmentation for crop and weed detection using CNNs. In IEEE International Conference on Robotics and Automation (ICRA), Brisbane, Australia.

[6] You, A., Han, C., Choi, C., & Lee, D. (2020). DNN-based semantic segmentation for weed and crop detection. Computers and Electronics in Agriculture, 175, 105528.

[7] Intawong, K., Intajag, S., & Apornsuksakul, A. (2013). A pixel-based quality measure for segmentation algorithms. In Computer Analysis of Images and Patterns (pp. 308–315). Springer.

[8] Roy, N., Kundu, S., Bhowmik, T. (2025). Enhanced maize cultivation: IoT based precision agriculture system. In Proc. 8th International Conference on Electronics, Materials Engineering & Nano-Technology (IEMENTech), Kolkata, India, pp. 1–6. 10.1109/IEMENTech65115.2025.10959491

[9] Kouzana, H., Nakib, A., & Dogaz, N. (2017). Microscopic image segmentation using branch and bound and game theory. In Metaheuristics for Medicine and Biology (pp. 211–226). Springer.

[10] Qi, S., Mei, K., Huang, R. (2021). Attanet: attention-augmented network for fast and accurate scene parsing. Proceedings of the AAAI Conference on Artificial Intelligence, 35(3), 2567–2575. 10.1609/aaai.v35i3.16359

[11] Kundu, S., Mukhopadhyay, S., Talukdar, R., Kaplun, D., Voznesensky, A., Sarkar, R. (2025). Deep learning model for grading carcinoma with Gini-based feature selection and linear production-inspired feature fusion. Scientific Reports, vol. 15, Article 21225. 10.1038/s41598-025-00217-w

[12] Li, S., et al. (2022). Ndnet: spacewise multiscale representation learning via neighbor decoupling for real-time driving scene parsing. IEEE Transactions on Neural Networks and Learning Systems, 35(6), 7884–7898. 10.1109/tnnls.2022.3221745

[13] Peng, J., et al. (2022). Pp-liteseg: a superior real-time semantic segmentation model. arXiv preprint arXiv:2204.02681.

[14] Pan, H., et al. (2022). Deep dual-resolution networks for real-time and accurate semantic segmentation of traffic scenes. IEEE Transactions on Intelligent Transportation Systems, 24(3), 3448–3460. 10.1109/tits.2022.3228042

[15] Li, H., et al. (2019). Dfanet: deep feature aggregation for real-time semantic segmentation. In Proceedings of the IEEE/CVF Conference on Computer Vision and Pattern Recognition (pp. 9522–9531).

[16] Kar, S., Mukhopadhyay, S., Kundu, S., Jha, D., Mallipeddi, R. (2026). TransE2UNet: Edge guided TransEfficientUNET for generalized colon polyp segmentation from endoscopy images. In Medical Image Understanding and Analysis, Lecture Notes in Computer Science, vol. 15918, pp. 3–16. Springer Nature Switzerland. 10.1007/978-3-031-98694-9_1

[17] Yu, C., et al. (2018). Bisenet: bilateral segmentation network for real-time semantic segmentation. In Proceedings of the European Conference on Computer Vision (ECCV) (pp. 325–341).

[18] Yu, C., et al. (2021). Bisenet v2: bilateral network with guided aggregation for real-time semantic segmentation. International Journal of Computer Vision, 129(11), 3051–3068. 10.1007/s11263-021-01515-2

[19] Kundu, S., Roy, N., Talukdar, R., Das, S., Mukhopadhyay, S., Basu Mallik, B. (2025). Classification and identification of potato leaf disease using ensembling of attention-aided deep neural networks through feature fusion. In Proc. 9^th^ International Conference on Information Technology (InCIT), Phuket, Thailand, pp. 499–505. 10.1109/InCIT66780.2025.11276115

[20] Shijie, H., et al. (2022). Real-time semantic segmentation via spatial-detail guided context propagation. IEEE Transactions on Neural Networks and Learning Systems.

[21] Poudel, R.P.K., Liwicki, S., Cipolla, R. (2019). Fast-scnn: fast semantic segmentation network. arXiv preprint arXiv:1902.04502.

[22] Kundu, S., Mukhopadhyay, S., Talukdar, R., Das, S., Adhikari, S. (2025). JUST: Towards jute pest classification by combination of supervised learning and triplet loss aided contrastive learning. Iran Journal of Computer Science, 8(2), 365–378. 10.1007/s42044-024-00222-8

[23] Kundu, S., Roy, N., Talukdar, R., Das, S., Mukhopadhyay, S., Basu Mallik, B. (2024). RAF2Net: Automated grading of renal cell carcinoma utilizing attention-enhanced deep learning models through feature fusion.

[24] Kundu, S., Talukdar, R., Roy, N., Das, S., Basu, S., Mukhopadhyay, S. (2024). Majority-LCL: Towards malaria cell detection using label contrastive learning and majority voting ensembling.

[25] Kundu, S., Roy, N., Chatterjee, P., Mitra, D., Bhowmik, T. (2025). Automated detection of rice crop disorder using deep learning techniques. In Pattern Recognition. ICPR 2024 International Workshops and Challenges, pp. 154–166. Springer Nature Switzerland. 10.1007/978-3-031-87657-8_11

[26] Kundu, S., Mukhopadhyay, S., Das, S., Mallipeddi, R. (2025). Reward: Renal carcinoma grade prediction using weighted confidence-based ensembling of attention integrated deep neural networks. SSRN Electronic Journal. 10.2139/ssrn.5107551

[27] Kundu, S., Talukdar, R., Das, S., Mukhopadhyay, S., Mallick, S., Basu Mallik, B. (2025). Fuzzy-CL: Fuzzy rank-based ensembling aided contrastive learning for malaria detection using red blood cell smears. In Proc. International Conference on Communication, Computing, Networking, and Control in Cyber-Physical Systems (CCNCPS), Dubai, UAE, pp. 325–330. 10.1109/CCNCPS66785.2025.11135766

[28] Mukherjee, A., Roy, N., Kundu, S., Panja, A., Chatterjee, S. (2025). EdgeSwarm: Edge-enabled opportunistic UAV network-assisted robotic swarm for postdisaster relief. In Recent Advances in Artificial Intelligence and Smart Applications, Lecture Notes in Networks and Systems, vol. 1380, pp. 167–179.Singapore. 10.1007/978-981-96-5822-0_15

